# The *Salmonella* pathogenicity island 1-encoded small RNA InvR mediates post-transcriptional feedback control of the activator HilA in *Salmonella*

**DOI:** 10.1101/2024.11.21.624761

**Authors:** Yutong Hou, Kyungsub Kim, Fatih Cakar, Yekaterina A. Golubeva, James M. Slauch, Carin K. Vanderpool

**Affiliations:** Department of Microbiology, University of Illinois at Urbana-Champaign, Urbana, Illinois, USA

**Author notes:** Department of Medicine, Division of Infectious Diseases, Massachusetts General Hospital, Massachusetts, USA. Regenerative and Restorative Medicine Research Center (REMER), Research Institute for Health Sciences and Technologies (SABITA), Istanbul Medipol University, Istanbul, Turkey (Fatih Cakar). Address correspondence to Carin Vanderpool.

## Abstract

*Salmonella* Pathogenicity Island 1 (SPI1) encodes a type three secretion system (T3SS) essential for *Salmonella* invasion of intestinal epithelial cells. Many environmental and regulatory signals control SPI1 gene expression, but in most cases, the molecular mechanisms remain unclear. Many of these regulatory signals control SPI1 at a post-transcriptional level and we have identified a number of small RNAs (sRNAs) that control the SPI1 regulatory circuit. The transcriptional regulator HilA activates expression of the genes encoding the SPI1 T3SS structural and primary effector proteins. Transcription of *hilA* is controlled by the AraC-like proteins HilD, HilC, and RtsA. The *hilA* mRNA 5’ untranslated region (UTR) is ~350-nuclotides in length and binds the RNA chaperone Hfq, suggesting it is a likely target for sRNA-mediated regulation. We used the rGRIL-seq (reverse global sRNA target identification by ligation and sequencing) method to identify sRNAs that bind to the *hilA* 5’ UTR. The rGRIL-seq data, along with genetic analyses, demonstrate that the SPI1-encoded sRNA InvR base pairs at a site overlapping the *hilA* ribosome binding site. HilD and HilC activate both *invR* and *hilA*. InvR in turn negatively regulates the translation of the *hilA* mRNA. Thus, the SPI1-encoded sRNA InvR acts as a negative feedback regulator of SPI1 expression. Our results suggest that InvR acts to fine-tune SPI1 expression and prevent overactivation of *hilA* expression, highlighting the complexity of sRNA regulatory inputs controlling SPI1 and *Salmonella* virulence.

**IMPORTANCE:** *Salmonella* Typhimurium infections pose a significant public health concern, leading to illnesses that range from mild gastroenteritis to severe systemic infection. Infection is initiated and requires a complex apparatus that the bacterium uses to invade the intestinal epithelium. Understanding how *Salmonella* regulates this system is essential for addressing these infections effectively. Here we show that the small RNA (sRNA) InvR imposes negative feedback regulation on expression of the invasion system. This work underscores the role of sRNAs in *Salmonella’s* complex regulatory network, offering new insights into how these molecules contribute to bacterial adaptation and pathogenesis.

## INTRODUCTION

*Salmonella* serovars are enteric foodborne pathogens that infect humans and animals by interacting with intestinal tissue and triggering gastrointestinal disease (1, 2). Upon entering the small intestine, numerous environmental signals trigger expression of a Type 3 Secretion System (T3SS) encoded by *Salmonella* Pathogenicity Island 1 (SPI1) (3–7). The bacterium uses the T3SS to inject effector proteins from the bacterial cytoplasm into the host cell, triggering uptake by the non-phagocytic epithelial cells and promoting inflammatory diarrhea (8–10). *Salmonella* cells that cross the gut epithelium are engulfed by macrophages via phagocytosis. The bacteria can then replicate in macrophages, leading to a potentially lethal systemic infection (11, 12).

*Salmonella* possesses complex regulatory systems that translate environmental signals into transcriptional and translational regulation of SPI1 to allow invasion at the appropriate time and place in the host. The SPI1 locus encodes HilA, the transcriptional activator of the SPI1 T3SS structural genes and primary effector proteins (13, 14). At the transcriptional level, *hilA* is regulated by a complex feedforward loop consisting of three AraC-like transcriptional regulators, HilD, HilC, and RtsA (Fig. 1A). These three factors autoregulate their own transcription and regulate one another’s transcription, along with transcription of *hilA* (15–18). HilD is the dominant regulator and the integrator of upstream signals (6). Loss of either *hilD* or *hilA* significantly reduces *Salmonella* intestinal colonization and internalization (18). Previous genetic analyses demonstrate that multiple regulatory factors or environmental cues modulate *hilA* and *hilD* expression (6, 19–21). Many of these regulators alter *hilD* expression at the post-transcriptional level, thereby indirectly regulating *hilA* (22–24). These observations encouraged us to identify additional regulatory factors acting at the post-transcriptional level, such as small RNAs (sRNAs).

**Figure 1.**
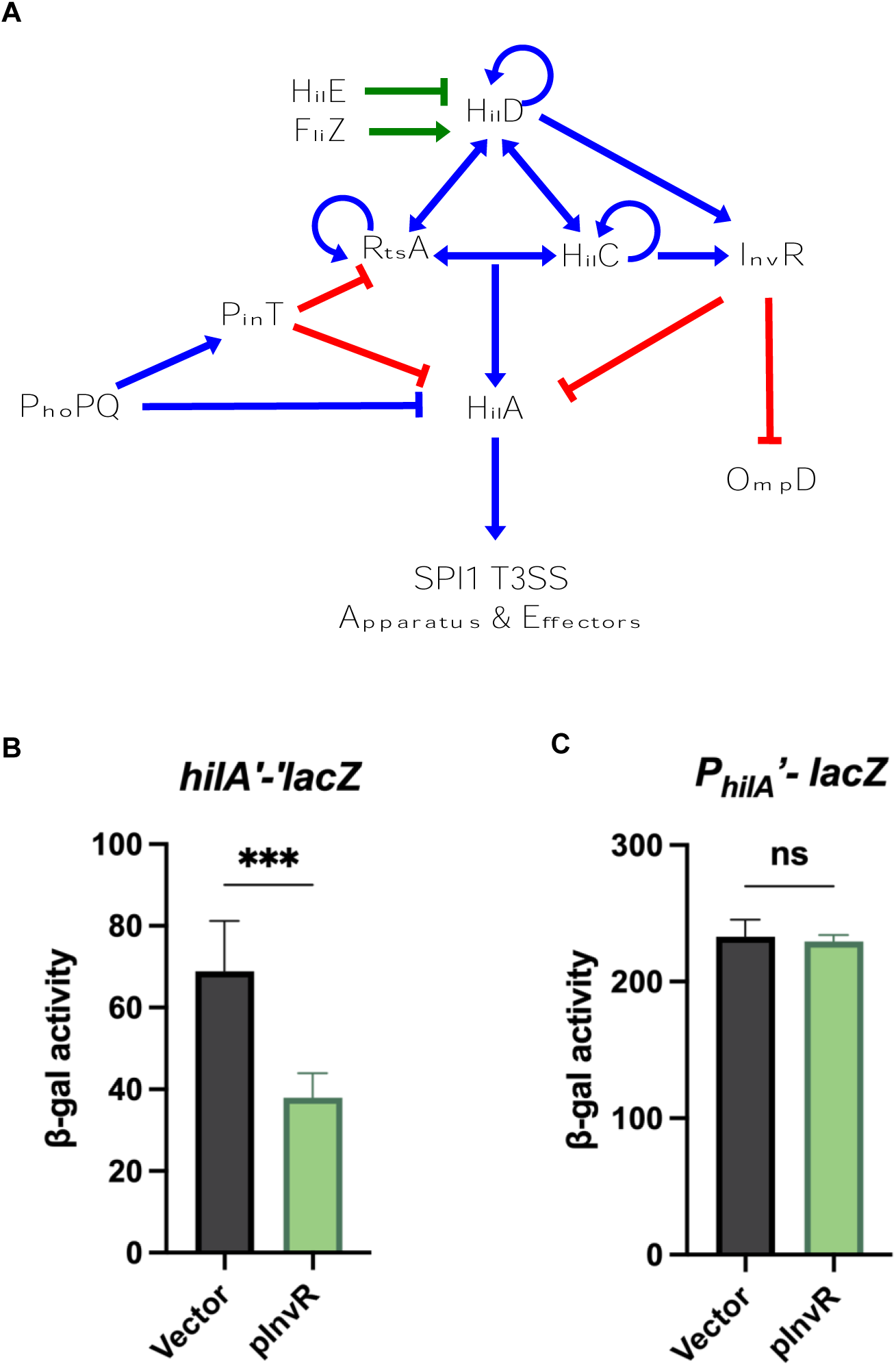
Regulation model and analysis of InvR-mediated control of SPI1 T3SS. A. Simplified regulatory model of the SPI1 T3SS and related regulators. Blue lines indicate transcriptional regulation, green lines indicate regulation at the protein level, red lines indicate regulation at the posttranscriptional level. β-galactosidase activity in *Salmonella* strains containing the (B) *hilAʹ-ʹ lacZ* translational and (C) P*_hilA_ʹ-lacZ* transcriptional fusions. Strains carrying vector control or InvR expression (pInvR) plasmids were grown under SPI-1-inducing conditions. β-galactosidase activity is presented as means ± standard deviations. Error bars represent the standard deviations from three independent experiments, analyzed using an unpaired t-test (n = 3). Statistical significance is indicated: *P < 0.05; **P < 0.005; ***P < 0.0005; ns, not significant. Strains used: JS2333, JS2217 with indicated plasmids.

Bacterial sRNAs are typically non-coding RNAs that exert post-transcriptional regulation by base pairing with target mRNAs to impact their stability, transcription elongation, or translation (25–27). Most sRNAs are transcribed from independent loci or processed from mRNA 3’ untranslated regions (UTRs) (28). Numerous sRNAs target mRNA 5ʹ UTRs and can work through various mechanisms (29). Many sRNAs in enteric bacteria require the RNA-binding protein Hfq, a homo-hexameric RNA chaperone that binds sRNAs and their target mRNAs to promote specific sRNA-mRNA pairing (30–33). Base pairing interactions can result in occlusion of the mRNA translation initiation region, which blocks the access of the 30S ribosome, hence inhibiting translation (32). Moreover, sRNA binding can initiate mRNA turnover mediated by the ribonuclease RNase E degradosome (34–36). Small RNAs like DsrA, RprA and ArcZ (37), as well as RydC and CpxQ (38) can also alter the transcription elongation of their targets by modulating the efficiency of Rho-dependent termination.

Our previous studies in *Salmonella* have revealed that SPI1 regulators are common targets of sRNAs responding to a variety of conditions. For example, FnrS and ArcZ are two sRNAs that respond to oxygen tension (22). FnrS is activated by Fnr under anaerobic conditions, whereas ArcZ is expressed under aerobic conditions (35, 39, 40). Both FnrS and ArcZ inhibit *hilD* translation by base-pairing with the 5’ UTR of *hilD* mRNA. MicC was also shown to repress *hilD* translation by base-pairing at 5’ UTR of *hilD* mRNA (23). Moreover, under the low Mg^2+^, low pH conditions of the phagosome, PhoPQ activates the sRNA PinT, which represses both *hilA* and *rtsA* translation by directly base-pairing within the 5’ UTRs near the ribosome binding sites (41). More recent studies have uncovered two sRNAs, SdsR and Spot42, that bind at the unusual 300-nt 3’ UTR of *hilD* to control mRNA stability (24, 42). Altogether, these results provide increasing evidence that sRNAs add an extra layer of regulation to SPI1, assuring quick adaptation in different environmental niches. Given that the *hilA* mRNA has a 350-nt long 5’ UTR, we hypothesized that additional sRNAs regulate SPI1 through interaction with the *hilA* mRNA.

InvR (<u>inv</u>asion gene-associated <u>R</u>NA) is an 87-nt long sRNA encoded by the *invR* gene located at one end of SPI1. InvR is an Hfq-binding sRNA (43). Prior to this work, the only known target of InvR was the *ompD* mRNA, which encodes an outer membrane porin. InvR represses the translation of *ompD* by direct base pairing just downstream of the translation start site (43). Although *invR* is located within the SPI1 locus, it was not previously shown to be associated with SPI1 regulation. Here, we demonstrate that InvR regulates production of the SPI1 transcriptional activator HilA. Using computational analysis and genetic approaches, we revealed that the molecular mechanism of InvR-mediated regulation of *hilA* is direct translational repression. We show that InvR serves as a feedback inhibitor between HilD and HilA in virulence-relevant conditions. This work broadens the sRNA-mediated post-transcriptional regulatory network of SPI1 under virulence conditions and provides new insight into how sRNAs fine-tune gene expression in bacteria during complex processes like virulence.

## RESULTS

### Identification of sRNAs that regulate SPI1 through the *hilA* 5’ UTR

To investigate potential sRNA regulators of SPI1 that act through the 350-nt *hilA* 5’ UTR in *Salmonella*, we utilized several bioinformatic tools including IntaRNA (44–46), TargetRNA (47), and Starpicker (48). We also performed rGRIL-Seq (reverse Global sRNA Target Identification by Ligation and Sequencing (49)) to capture sRNAs that specifically bind the *hilA* 5’ UTR under SPI1-inducing conditions. In this experiment, the *hilA* 5’ UTR plus 111 nt of *hilA* coding sequence was ectopically produced in *Salmonella* with co-production of T4 RNA ligase. Chimeric RNAs containing portions of the *hilA* 5’ UTR represent putative sRNA-*hilA* mRNA pairs that were in close proximity (e.g., as base paired complexes on Hfq) within the cell. These chimeric RNAs were enriched using bead-immobilized complementary oligonucleotides (see details in Materials and Methods). The rGRIL-seq results (Fig. S1) identified two predominant sRNAs that interact with the *hilA* mRNA. Many of the chimeric RNAs contained the region corresponding to approximately +15 to +25 nt of the *hilA* mRNA (relative to the start codon) and +9 to +24 nt of the PinT sRNA (relative to the transcription start site), indicating PinT binds in the early coding region of *hilA* mRNA. These results are consistent with our published results characterizing PinT-*hilA* mRNA interactions (41). In addition, we identified numerous *hilA* mRNA-InvR chimeric RNAs (Fig. S1B) containing a region of *hilA* mRNA from −14 to −21 nt (relative to the start codon) and +21 to +33 nt of the InvR sRNA (relative to the transcription start site). A similar *hilA* mRNA-InvR RNA chimera pattern was also captured (but not characterized) in a recently published iRIL-Seq paper (24). These data suggested that the sRNA InvR is also a direct regulator of the *hilA* mRNA.

### InvR is a direct negative regulator of *hilA* translation

To study InvR and *hilA* mRNA interactions, we tested whether ectopic production of InvR affected *hilA* transcription or translation. We cloned *invR* from *Salmonella* Typhimurium strain 14028s on a plasmid under the control of an IPTG-inducible promoter (50, 51) and introduced the plasmid into *Salmonella* strains containing either *hilA’-’lacZ* translational fusion or *hilA* promoter transcriptional fusion. Ectopic production of InvR reduced fusion activity by about 50% only for the *hilA* translational fusion but not the *hilA* promoter transcriptional fusion (Fig. 1B and C).

Our data suggest that InvR directly represses *hilA* by base pairing in the *hilA* 5’ UTR. However, based on the feedforward loop model of SPI1 regulation, we reasoned that InvR could also be affecting an upstream SPI1 regulator (18, 52). The sRNA PinT, for example, regulates translation of both *hilA* and *rtsA* (41). We tested the effect of InvR on other SPI1 regulators using translational fusions. Ectopic production of InvR had no significant impact on the *hilDʹ-ʹlacZ* translational fusion in *Salmonella* under SPI1-inducing conditions (Fig. 2A). Overproduction of InvR slightly increased levels of *rtsAʹ-ʹlacZ* in *Salmonella* (Fig. 2B), but the deletion of *invR* did not impact *rtsAʹ*-*ʹlacZ* fusion activity (Fig. 2C). Similarly, the overproduction of InvR slightly decreased the translation levels of *hilCʹ*-*ʹlacZ* in *Salmonella* (Fig. 2D), but again, deletion of *invR* had no effect on *hilC* translation under SPI-1-inducing conditions (Fig. 2E).

**Figure 2.**
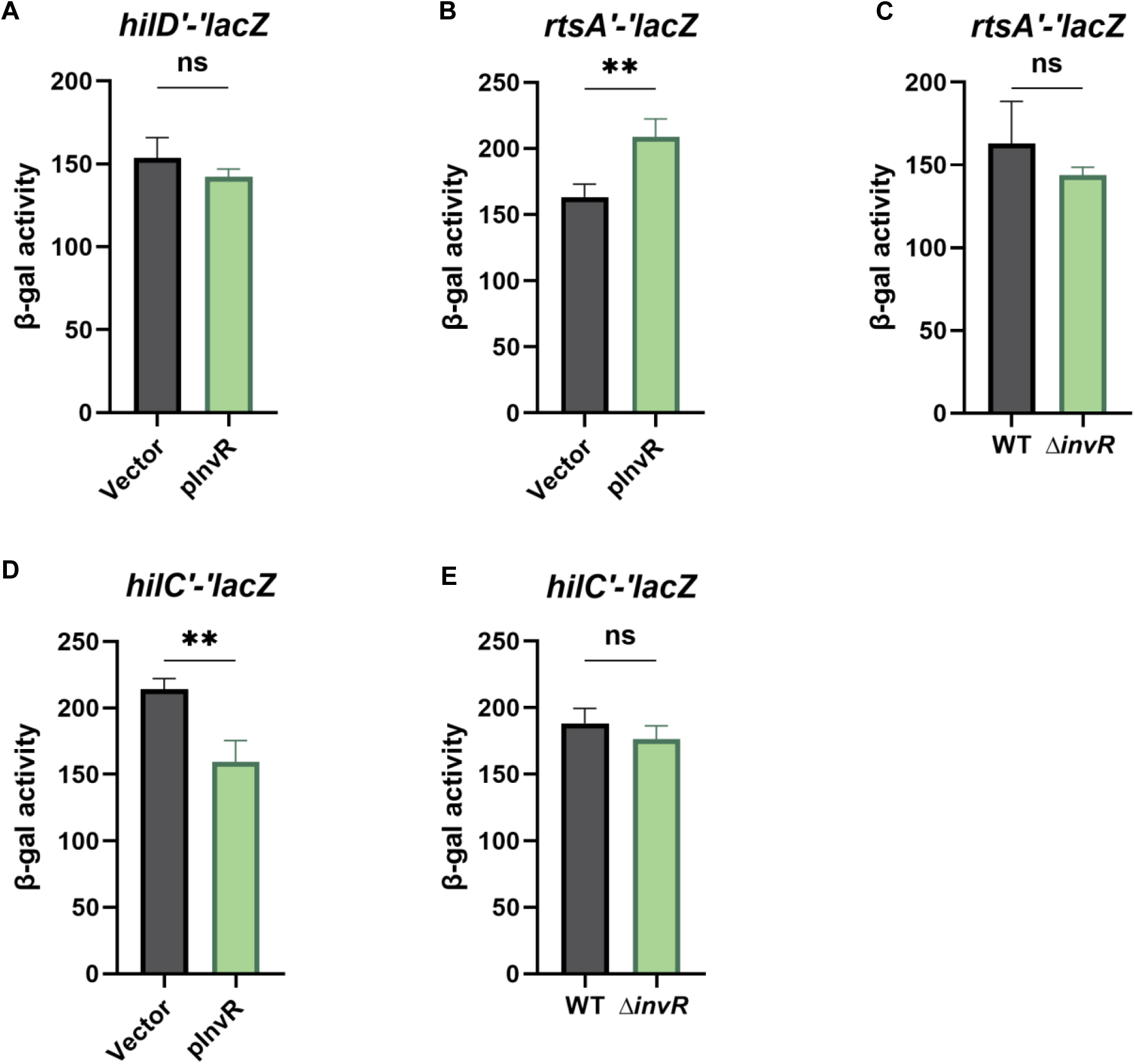
InvR does not regulate *hilD, rtsA,* or *hilC* translation. β-galactosidase activity in *Salmonella* strains containing translational fusions: (A) *hilDʹ-ʹlacZ,* (B) *rtsAʹ-ʹlacZ,* or (D) *hilCʹ-ʹlacZ* with vector control or InvR expression (pInvR) plasmids. Strains were grown under SPI-1-inducing conditions. C. β-galactosidase activity in *Salmonella* strains containing a *rtsAʹ-ʹlacZ* translational fusion in wild-type or Δ*invR* background. Strains were grown under SPI-1-inducing conditions. E. β-galactosidase activity in *Salmonella* strains containing a *hilCʹ-ʹlacZ* translational fusion in wild-type or Δ*invR* background. Strains were grown under SPI-1-inducing conditions. control or InvR expression (pInvR) plasmids were grown under SPI-1-inducing conditions. β-galactosidase is presented as means ± standard deviations. Error bars represent the standard deviations from three independent experiments, analyzed using an unpaired t-test (n = 3). Statistical significance is indicated: *P < 0.05; **P < 0.005; ***P < 0.0005; ns, not significant. Strains used: JS892, JS2334, JS2675, JS2676, JS2677 with indicated plasmids.

To further investigate the regulation of *hilA*, we used a *hilA*ʹ-ʹ*lacZ* translational fusion in *E. coli* under the control of the P_BAD_ promoter (arabinose-inducible). This eliminates any requirement for *Salmonella-*specific transcription signals. Ectopic production of InvR reduced *hilA* translation level by ~58% in P_BAD_*-hilA’-’lacZ* (Fig. 3A) fusions. Since InvR was reported to repress *ompD* translation (43), we used a P_BAD_*-ompD’-’lacZ* translational fusion as a positive control and a P_BAD_*-hilD’-’lacZ* translational fusion as a negative control. Ectopic production of InvR repressed P_BAD_-*ompD*’-’*lacZ* (Fig. 3B) but did not affect P_BAD_-*hilD’-’lacZ* (Fig. 3C) fusions. Because InvR represses a *hilA* fusion in the absence of any other SPI1 components (in *E. coli*) and we see no evidence of significant InvR regulation of any other SPI1 regulator or *hilA* promoter activity, we concluded that InvR affects *hilA* directly, likely by base pairing.

**Figure 3.**
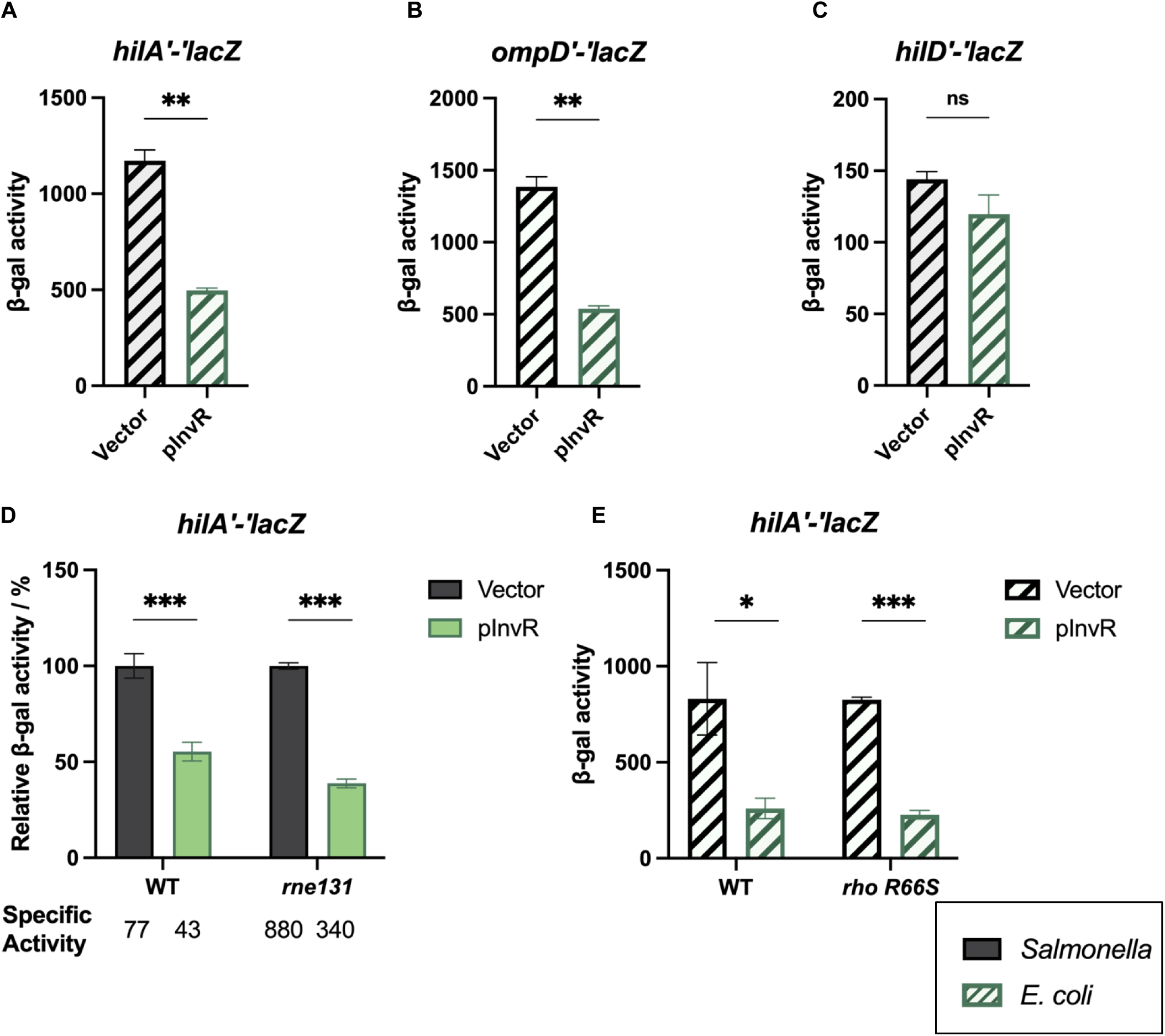
InvR represses *hilA* translation independent of Rho and RNase E. β-galactosidase activity in *E. coli* containing translational fusions: (A) *hilAʹ-ʹlacZ,* (B) *ompDʹ-ʹlacZ,* (C) *hilDʹ-ʹlacZ* with vector control or InvR expression (pInvR) plasmids. Strains were grown as described in Materials and Methods. D. β-galactosidase activity in *Salmonella* strains containing the *hilAʹ-ʹlacZ* translational fusion in WT (*rne^+^*) or *rne131* backgrounds. Strains carrying vector control or InvR expression (pInvR) plasmids were grown under SPI-1-inducing conditions. E. β-galactosidase activity in *E.coli* strains containing *hilAʹ-ʹlacZ* translational fusion in wild type (WT) background or *rho-*R66S background. Strains carrying vector control or InvR expression (pInvR) plasmids were grown as described in Materials and Methods. control or InvR expression (pInvR) plasmids were grown under SPI-1-inducing conditions. Relative β-galactosidase units were calculated by normalizing β-galactosidase activity to that of the wild-type strain with vector control and are presented as means ± standard deviations. Error bars represent the standard deviations from three independent experiments, analyzed using an unpaired t-test (n = 3). Statistical significance is indicated: *P < 0.05; **P < 0.005; ***P < 0.0005; ns, not significant. Strains used: JMS6505, GH05, JMS6500, JS2678, GH08 with indicated plasmids.

InvR base pairing within the long *hilA* 5’ UTR could promote RNase E-dependent mRNA degradation (53–55) or premature Rho-dependent termination (37, 38) To test whether the RNA degradosome is involved in InvR-mediated regulation of *hilA*, we introduced the *rne131* allele (34) into the *Salmonella hilA’-’lacZ* reporter strain. Although *hilA* fusion activity levels were significantly increased in the *rne131* background compared to the wild-type *rne*^+^ strain (Fig. 3D, see specific activity), we observed that InvR-mediated repression occurred to the same extent in both backgrounds. To test whether Rho-dependent transcription termination impacted InvR-mediated repression, we examined *hilA’-’lacZ* activity in the *E. coli* fusion strain after the introduction of the termination-defective *rho-*R66S allele (38, 56). There was no Rho-dependent difference in the basal (vector control) level of activity of the *hilA’-’lacZ* fusion, suggesting that the *hilA* 5’ UTR is not a substrate for Rho-dependent termination. Moreover, InvR still repressed *hilA* translation in the *rho-*R66S background (Fig. 3E). These data suggest that neither the RNA degradosome nor Rho factor play any important role in InvR-mediated regulation of *hilA.* Thus, we hypothesized that InvR binds to the *hilA* 5’ UTR to directly repress translation.

### Minimal region of *hilA* mRNA 5’ UTR required for InvR-mediated repression

To define the region of the *hilA* mRNA required for InvR-mediated repression, we created a series of translational fusion constructs in *E. coli* corresponding to successive 5’ deletions from the transcription start site of *hilA* mRNA (Fig. 4A). We tested the effect of InvR ectopic production on each of the fusions. Although the deletions in the *hilA* 5’ UTR had minor impacts on the basal level of fusion activity, fusions L1 (−285 to +30) through L8 (−40 to +30) were all still repressed by InvR (Fig. 4B). These data suggest that most of the *hilA* 5’ UTR is dispensable for regulation by InvR and that sequences from −40 to +30 (relative to the *hilA* start codon) confer InvR-dependent regulation. The S fusion, which contains −30 to +30 relative to the *hilA* start codon (Fig. 4B) was not regulated by InvR, suggesting that sequences important for regulation are missing from this fusion.

**Figure 4.**
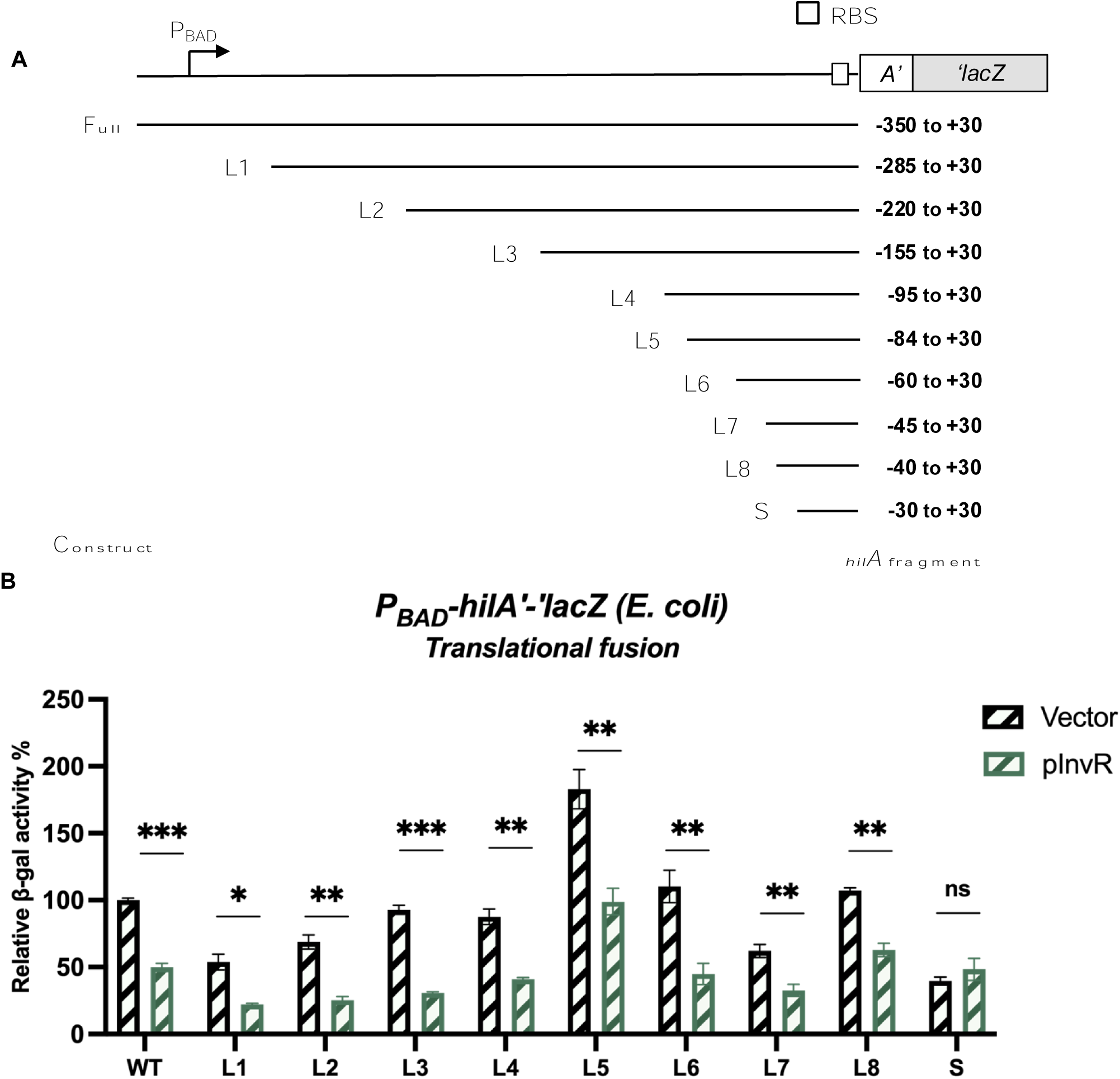
Region of the *hilA* 5’UTR required for InvR-mediated repression. A. Schematic representation of the truncated *hilA* 5’UTR in translational fusions. Fragments of *hilA* 5’UTR were deleted sequentially from the 5’ end of *hilA* between the transcription start site and 30 nt upstream of AUG start codon (−30). The +30 site (relative to the start codon) of *hilA* was fused to *lacZ* to create translational fusions. B. β-galactosidase activity in *E. coli* containing different truncation mutants of *hilAʹ-ʹlacZ*. Strains carrying vector control or pInvR expression plasmids were grown as described in Materials and Methods. control or InvR expression (pInvR) plasmids were grown under SPI-1-inducing conditions. Relative β-galactosidase units were calculated by normalizing β-galactosidase activity to that of the wild-type strain with vector control and are presented as means ± standard deviations. Error bars represent the standard deviations from three independent experiments, analyzed using an unpaired t-test (n = 3). Statistical significance is indicated: *P < 0.05; **P < 0.005; ***P < 0.0005; ns, not significant. Strains used: JMS6505, GH407, GH408, GH14, GH409, GH663, GH587, GH666, GH667, GH589, GH576 with indicated plasmids.

### InvR represses *hilA* by direct binding with the ribosome binding site of *hilA* mRNA

To further define the InvR binding site on *hilA* mRNA, we combined evidence from our rGRIL-seq data (Fig. S1), deletion analysis (Fig. 4) and binding site prediction by the IntaRNA program (46). These three lines of evidence suggest that InvR base pairs with sequences at the ribosome binding site (RBS) of *hilA* mRNA (Fig. 5A). Interestingly, the region of InvR predicted to base pair with *hilA* differs from that known to base pair with the *ompD* mRNA (Fig. 5B; (43)). To investigate whether this predicted base pairing is required for the InvR-mediated regulation of *hilA*, we introduced six different mutations into *invR* that should disrupt the *hilA* mRNA-InvR base pairing interactions, but not impact InvR base pairing with the *ompD* mRNA. All six InvR mutants lost the ability to repress the wild-type *hilA’-’lacZ* fusion in *E. coli* (Fig. 5C) but retained the ability to repress *ompD* translation at the same levels as wild-type InvR (Fig. 5D). These results show that the mutations do not influence InvR structure or stability, but specifically impair regulation of *hilA,* consistent with the base pairing prediction. Next, we introduced a compensatory mutation in *hilA* (denoted as *hilA-mut1*) that should disrupt the base pairing interaction with wild-type InvR and restore the interaction with InvR-mut1. We note that the *hilA*-mut1 mutation reduced translation of the mutant fusion to ~30% of the level of the wild-type fusion (Fig. 5E), likely due to the proximity to the RBS. Moreover, this mutation in *hilA* did not disrupt wild-type InvR-mediated regulation. However, consistent with our prediction, the regulation by the InvR-mut1 allele was restored by the compensatory mutation and reduced the translation of *hilA-mut* by ~71%. These data are consistent with the model that InvR base pairs with sequences overlapping the RBS of *hilA* mRNA to directly repress *hilA* translation.

**Figure 5.**
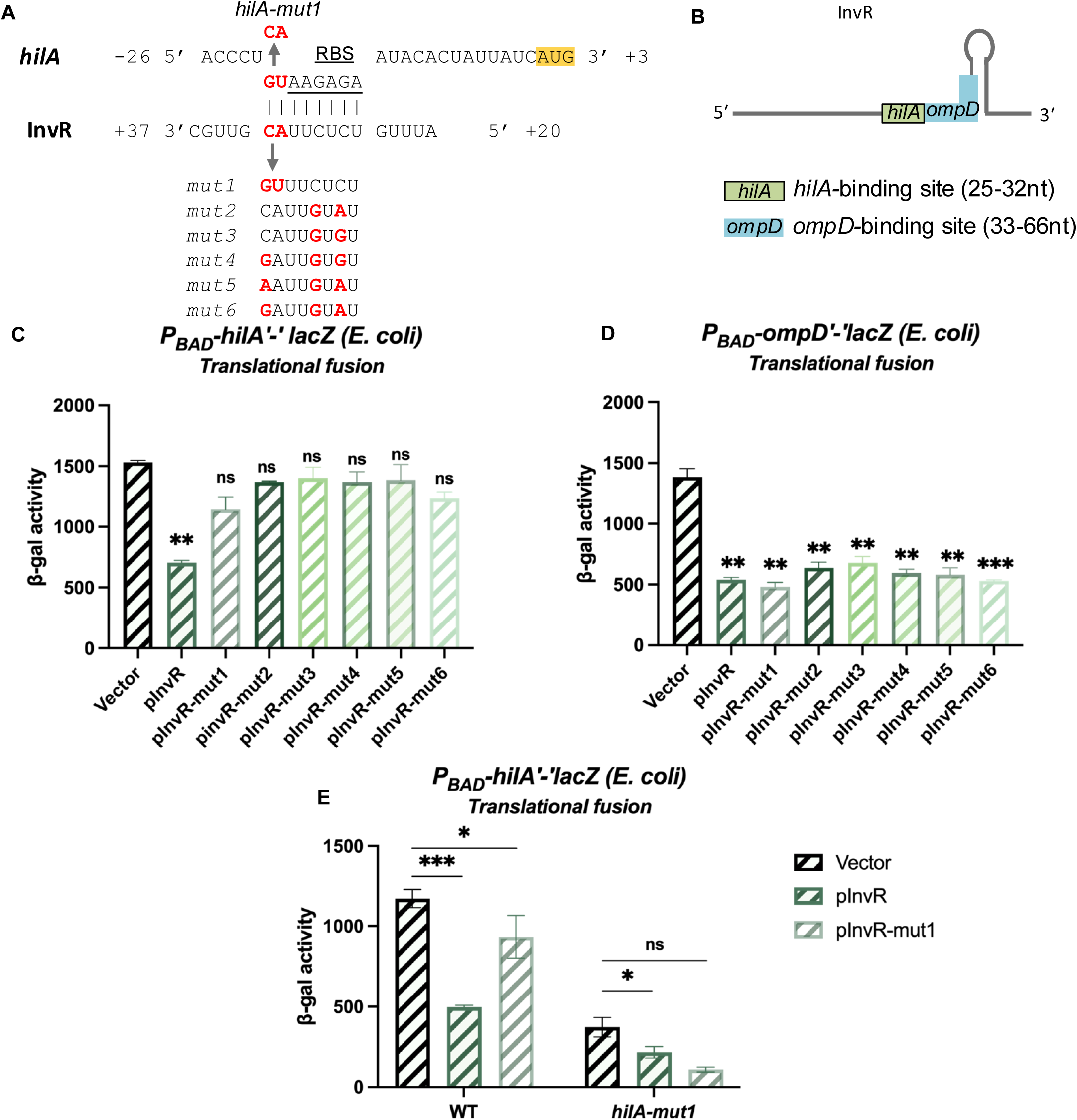
InvR-mediated *hilA* repression involves base pairing at the *hilA* ribosome binding site. A. Predicted base-pairing interactions between InvR and *hilA* mRNA. For *hilA*, nucleotides are numbered from the translational start site. Red font indicates nucleotides where mutations were created in P_BAD_-*hilAʹ-ʹlacZ* or pInvR. B. Schematic of InvR sRNA seed regions for *hilA* or *ompD* regulation. C. and D. β-galactosidase activity in *E. coli* strains containing the wild type *hilAʹ-ʹlacZ* translational or *ompDʹ-ʹlacZ* translational fusions; strains carry vector control, wild-type (pInvR) or mutant (pInvR-mut1 through mut6) plasmids. E. β-galactosidase activity in *E. coli* strains containing the wild type or mutant *hilAʹ-ʹlacZ* translational fusions; strains carry the wild type (pInvR) or mutant (pInvR-mut1) expression plasmids. β-galactosidase activity is presented as means ± standard deviations. Error bars represent the standard deviations from three independent experiments, analyzed using an unpaired t-test (n = 3). Statistical significance is indicated: *P < 0.05; **P < 0.005; ***P < 0.0005; ns, not significant. Strains used: GH02, GH05 with indicated plasmids.

### Potential feedback regulation by HilD to HilA through InvR

HilD was reported to control *invR* transcription (43, 57). We confirmed HilD-dependent *invR* transcription using an *invR’-lacZ*^+^ transcriptional fusion. Activity of the *invR’-lacZ*^+^ fusion in a Δ*hilD* mutant background was reduced to 17% of the activity in the wild-type background (Fig. 6A). To determine if HilA has any effect on *invR* transcription, we compared expression of the fusion in a Δ*hilA* background under SPI1-inducing conditions. We observed that deletion of *hilA* resulted in a 2-fold upregulation of *invR* transcription. Our previous work revealed that transcription of *hilD* is controlled by long-distance effects of H-NS binding and that deletion of *hilA* increases *hilD* transcription (58). Thus, we conclude that higher levels of *hilD* transcription in the Δ*hilA* background led to more HilD-dependent activation of *invR*.

**Figure 6.**
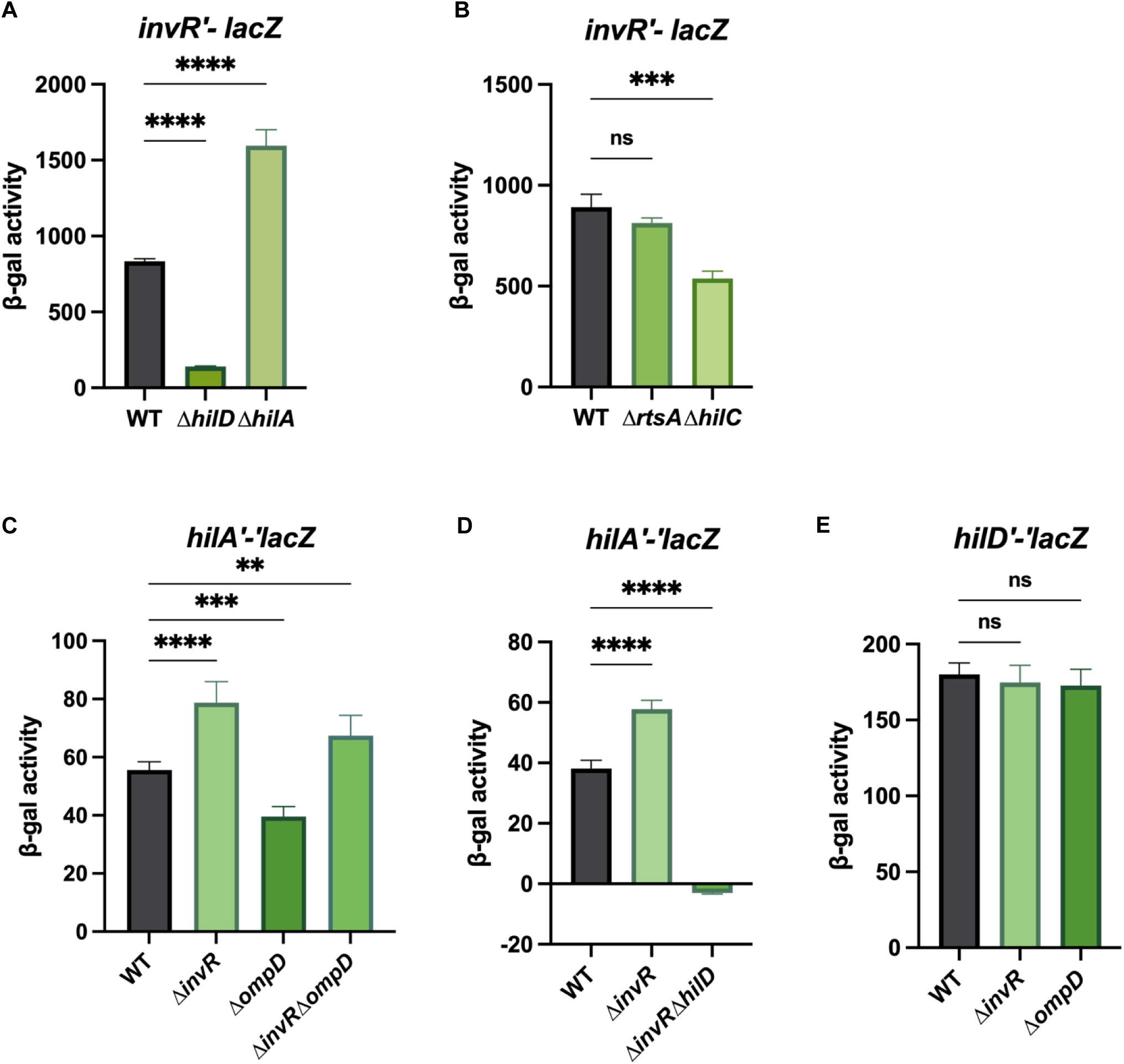
Feedback inhibition of InvR on *hilA* translation. A. β-galactosidase activity in *Salmonella* strains containing an *invRʹ-lacZ^+^* transcriptional fusion in wild-type, Δ*hilD* or Δ*hilA* background. B. β-galactosidase activity in *Salmonella* strains containing an *invRʹ-lacZ^+^* transcriptional fusion in wild-type, Δ*rtsA* or Δ*hilC* background. C. β-galactosidase activity in *Salmonella* strains containing *hilAʹ-ʹlacZ* translational fusion in wild-type, Δ*invR*, Δ*ompD* or *ΔinvRΔompD* backgrounds. D. β-galactosidase activity in *Salmonella* strains containing *hilAʹ-ʹlacZ* translational fusion in wild-type, Δ*invR*, Δ*hilD* or *ΔinvRΔhilD* background. E. β-galactosidase activity in *Salmonella* strains containing *hilDʹ-ʹlacZ* translational fusion in wild-type, *ΔinvR* or *ΔompD* backgrounds. β-galactosidase activity is presented as means ± standard deviations. Error bars represent the standard deviations from three independent experiments, analyzed using an unpaired t-test (n = 3). Statistical significance is indicated: *P < 0.05; **P < 0.005; ***P < 0.0005; ns, not significant. Strains used: JS2679-2683, JS2333, JS2684-2687, JS892, JS2688-2689.

HilD, HilC, and RtsA can form homodimers and heterodimers and bind to essentially the same sequences to activate transcription (16, 59). Therefore, we also examined *invR* transcription levels in the absence of *rtsA* and *hilC* (Fig. 6B). Interestingly, *invR* transcriptional fusion activity in the *hilC* mutant background was ~42% of the levels in the wild-type background, showing that HilC contributes to *invR* activation. On the other hand, loss of RtsA had no significant effect on *invR* transcription.

Given that HilD activates both *hilA* and *invR* transcription, our model is that InvR represses *hilA* translation under SPI1-inducing or related conditions as a form of feedback control. We also hypothesized that InvR-mediated regulation of *hilA* might be impacted by the presence of InvR’s other mRNA target, *ompD*. To test this model, we created deletions of *invR* and *ompD* in the *hilA’-’lacZ* translational fusion strain. Deletion of *invR* increased *hilA’-’lacZ* activity under SPI1-inducing conditions (Fig. 6C), demonstrating that the presence of InvR limits *hilA* translation. Deletion of *ompD* reduced *hilA’-’lacZ* activity under SPI1-inducing conditions. The simplest explanation is that there is competition between the two InvR targets. In the absence of *ompD* mRNA, there is more InvR available to base pair with and repress *hilA* mRNA. As we expected, the *hilA* fusion activity in the *invR ompD* double deletion background was significantly increased compared to the wild type. Moreover, the *hilA* fusion showed almost no activity in the absence of HilD, regardless of the presence or absence of *invR* (Fig. 6D). The results are consistent with HilD being the dominant activator of both *hilA* and *invR*. On the other hand, the levels of *hilD’-’lacZ* were unchanged in *invR* and *ompD* mutant backgrounds compared to the wild type, showing that the impacts on *hilA* expression were not caused by changes in HilD levels (Fig. 6E). Thus, under SPI1 inducing conditions, both *invR* and *hilA* require HilD for their transcription. InvR represses *hilA* translation, providing feedback regulation between HilD and HilA (Fig. 1A).

### Impact of InvR on *Salmonella* virulence in mice

To examine the potential role of InvR during infection, we used competition assays in which mice were infected orally or intraperitoneally (IP). *Salmonella* virulence during oral infection depends on the SPI1 T3SS to allow invasion and systemic dissemination (8, 18, 60–62). Intraperitoneal infection bypasses the need for invasion and SPI1 is not required for infection by this route. We confirmed that when the Δ*invR* strain was co-cultured with the *invR^+^* strain in an otherwise wild-type background *in vitro*, both strains competed equally (data not shown). This suggests the absence of *invR* did not cause a generalized growth defect. When Δ*invR* and *invR^+^* strains were used to co-infect mice, we observed that they competed equally in both oral and IP infections after recovery from both the intestine and spleen (Table S1). These data suggest that the effects of InvR on SPI1 are too subtle to be detected in this assay.

## DISCUSSION

The central regulatory framework for the SPI1 T3SS is well understood at the transcriptional level. The AraC-like regulators HilD, HilC, and RtsA form a feedforward loop that activates the transcription of *hilA*, encoding the transcriptional activator of SPI1 structural genes (14, 18). Environmental cues are integrated primarily at the level of HilD (6). Here, we show that the SPI1-encoded sRNA InvR, transcriptionally induced by HilD and HilC, translationally represses *hilA*. InvR contributes to the feedback regulation of *hilA*, adding another layer of fine-tuning to the central regulatory network of SPI1.

Based on rGRIL-Seq data and mutational analyses, InvR base pairs at the ribosome binding site of the *hilA* mRNA to prevent translation initiation. This regulation is independent of RNase E and Rho (Fig. 2F and 2G). Pfeiffer, et al. (43) examined the effects of InvR on SPI1 secreted proteins and concluded that there was no significant effect, thus dismissing any role for InvR in SPI1 regulation. *Salmonella* RIL-seq also failed to pull down the InvR-*hilA* mRNA chimera (63). However, newly published iRIL-Seq data (64) did capture InvR-*hilA* mRNA chimeras. Both InvR and *hilA* mRNA fragments in the InvR-*hilA* mRNA chimeras are consistent with our rGRIL-Seq and mutagenesis data.

The previous characterization of InvR showed that the sRNA negatively regulates the translation of the outer membrane porin protein OmpD (43). The InvR-*ompD* mRNA interaction is mediated by InvR nucleotides 33-66 base pairing with a site in the *ompD* mRNA coding region. In contrast, a different region of InvR (nucleotides 25-32) base pairs with the *hilA* mRNA RBS region. The fact that the *hilA* mRNA seed region mutations in InvR did not interfere with InvR-mediated repression of *ompD* (Fig. 5D) proves that InvR uses distinct regions to interact with the two different target mRNAs. We noted that *hilA* expression was reduced in the absence of *ompD* but recovered in the Δ*invR ΔompD* double deletion background. This suggests that *hilA* and *ompD* mRNAs compete for InvR binding under SPI1-inducing conditions. The sRNA MicC also coordinately regulates SPI1 and outer membrane porin proteins. Transcriptionally induced by SlyA, MicC is a negative regulator of *hilD*, *ompC*, and *ompD* (23, 35). SPI1 expression is down-regulated in response to envelope stress, including problems with the beta-barrel assembly machinery, Bam (65). Coordinating SPI1 regulation and down-regulation of OmpD by InvR and MicC might lessen any stress in outer membrane assembly during infection. More studies are needed to understand these interconnections.

Despite these intriguing regulatory links, loss of InvR did not confer any significant changes in *Salmonella* fitness in our oral or systemic infection models. Expression of *hilA* increased only ~37% in the *ΔinvR* mutant compared to the wild-type strain (Fig. 6C). We reason that InvR-mediated post-transcriptional regulation of *hilA* might be too subtle in the overall context of SPI1 regulation to observe a strong phenotype in the animal model of infection. In contrast, several studies have shown a clear role for sRNA-mediated SPI1 regulation in *Salmonella* pathogenicity. For example, the Δ*micC* strain gained a fitness advantage during oral infection in a SPI-1-dependent manner (23). Conversely, the sRNAs SdsR and Spot42 both increase HilD production. Deletion of these sRNAs decreased *Salmonella* invasion in the mouse model, and this phenotype was dependent on SPI1 (24). Like InvR, loss of the other *hilA-*regulating sRNA PinT has no apparent effect on SPI-dependent intestinal invasion. However, the *pinT* mutant gained a virulence advantage during systemic infection, consistent with PinT’s role in coordinating SPI1 and SPI2 expression (41, 66). These findings demonstrated the crucial roles of sRNA in *Salmonella* virulence.

In this study, we demonstrated that HilD is essential for InvR production, consistent with previous work (43). We also showed that deletion of *hilC* reduced *invR* expression by ~50%, revealing HilC’s contribution to the regulation of InvR. HilD is required to induce and express *hilC* and *rtsA* (18). These observations are aligned with published HilD and HilC ChIP-Seq data (67), which captured HilD and HilC binding at the *invR* promoter region. In contrast, there was no significant effect of loss of RtsA on *invR* expression. HilD, HilC, and RtsA bind to the same sites in the *hilD*, *hilC*, *rtsA* and *hilA* promoters, although with slight differences in sequence recognition (16). Moreover, HilD, HilC and RtsA form both homodimers and heterodimers (59). It seems that there is a preference for HilD/HilC homodimers or heterodimers at the *invR* promoter.

While the SPI1 dominant activator, HilD, induces the transcription of InvR and *hilA*, InvR negatively regulates *hilA*. This type of feedback regulation is common in many biological systems and can be involved in various processes, including gene expression, signal transduction pathways, biosynthesis, and metabolism. It allows for precise control and fine-tuning of cellular responses. For example, the *Salmonella* flagellar regulatory network consists of several inter-connected feedback loops to maintain the dynamic control of flagellar assembly. While the master regulator FlhD_4_C_2_ activates FlgM (anti-sigma factor) and FliA (flagella-specific sigma factor), FlgM sequesters FliA by forming the FliA-FlgM protein complex, thus inhibiting FilA downstream function (68). Another interesting feedback regulation circuit is in the formation of biofilm. CsgD is the master transcriptional regulator of the structural proteins that form curli fibers. OmpR, as the response regulator in EnvZ/OmpR system, directly induces the transcription of CsgD as well as two sRNAs, OmrA and OmrB (69, 70). Intriguingly, these sRNAs use the same seed region to interact with *csgD* and *ompR* mRNA, repressing the translation initiation of two target mRNAs (70, 71). Thus, OmpR, CsgD, and OmrA/B comprise a complex feedback regulatory circuit to coordinate bacterial physiology and behavior.

In summary, the SPI1-encoded sRNA InvR negatively regulates HilA at the post-transcriptional level. Given that HilD and HilC control the transcription of *invR* and *hilA*, InvR creates a feedback loop. This regulatory setup combines transcriptional and posttranscriptional control, fine-tuning the expression of the SPI1 T3SS.

## MATERIALS AND METHODS

### Bacterial strains and plasmids

Bacterial strains and plasmids used in this study are listed in Tables S2 and S3. All *Salmonella* strains used in this study are isogenic derivatives of *Salmonella enterica* serovar Typhimurium strain 14028s (American Type Culture Collection [ATCC]). The reference genome assembly used in this study is from NCBI GenBank CP001361.1 and CP001363.2 (Assembly Accession: GCA_000022165, (72)). Chromosomal deletions and other mutations were made by λ red recombination (73) and moved into the appropriate strain background using P22 HT105/1 int-201 (P22)-mediated transduction (74). The transcriptional *lacZ* fusion to *invR* was made using FLP-mediated recombination with plasmid pKG137, as previously described (75).

Transcriptional *lacZ* fusions in *E. coli* were constructed by λ red recombination in the strain PM1805 as described previously (76). gBlocks (Integrated DNA Technologies) were used to construct strains containing *hilA*-mut1 mutants. PCR products containing truncated fragments of *hilA* mRNA (start site and end site are shown in Fig. 4A) were used to construct P_BAD_*-hilA*′*-*′*lacZ* truncated fusions in the strain PM1805 using λ Red recombination. The strain carrying the *rho-*R66S allele was made previously and transduced into appropriate strain background using P1 phage (56).

All oligonucleotide primers used in this study were synthesized by Integrated DNA Technologies and are listed in Table S4. PCR products were generated using Q5 Hot Start High-fidelity DNA polymerase (New England Biolabs, M0493S) or Phusion High-Fidelity DNA Polymerase (New England Biolabs, M0530S).

Plasmids encoding IPTG-inducible *invR* were constructed by amplifying *invR* sequence from strain 14028s using primers F-InvR_aatII/R-InvR_ecoRI (Table S2). The PCR products were cloned into a linearized and dephosphorylated pBRCS12 (also named as pBR-P_Llac_) Digested by AatII, EcoRI-HF, and rSAP; New England Biolabs, R0117S, R3101S, and M0371S). IntaRNA 2.0 was used to predict base pairing between InvR and *hilA* mRNA 5’ UTR (46). Plasmids containing *invR* mutants were constructed with Q5 Site-Directed Mutagenesis Kit (New England BioLabs, E0554S).

### Growth conditions and media

Strains were cultured in Lysogeny broth (LB; 10% Casein digest peptone, 5% yeast extract, 10% NaCl). For SPI1-inducing conditions, cells were grown in no-salt LB (NSLB; 10% tryptone, 5% yeast extract) overnight at 37°C with aeration, and then overnight grown cells were subcultured in high-salt LB (HSLB; 10% tryptone, 5% yeast extract, 10% NaCl). All strains were grown at 37°C, except for the strains containing the temperature-sensitive plasmids pCP20 or pKD46, which were grown at 30°C. When required, antibiotics were used at the following final concentrations: 100 μg/mL ampicillin (amp), 50 μg/mL kanamycin (kan), 10 μg/mL chloramphenicol (cm), 10 μg/mL tetracycline (tet).

### β-Galactosidase assays

β-Galactosidase assays were performed using a 96-well plate as previously described (77). Briefly, *Salmonella* strains were inoculated in NSLB medium and grown overnight at 37°C on a tissue culture rotator at 50 rpm (Fisher Scientific, 88-882-015). These cultures were then diluted 1:100 into 3 mL of HSLB medium in 13 mm tubes and grown statically at 37°C for 18 to 22 h. For *E. coli* cultures, strains were initially inoculated into 800 µL LB and grown overnight at 37°C with aeration, then subcultured 1% into 3 mL of LB medium with 100 μM isopropyl β-D-1-thiogalactopyranoside (IPTG) and 0.002% arabinose and grown at 37°C with aeration for 3 h. After incubation, cells were resuspended in 1.5 mL Z-buffer (60 mM Na_2_HPO_4_ ·7H_2_O, 40 mM NaH_2_PO_4_·H_2_O, 10 mM KCl, 1 mM MgSO_4_·H_2_O, pH 7.1) and OD_600_ was measured. Then, cells were permeabilized by adding 15µl 0.1% SDS and 20 µl chloroform. Nitrophenyl-β-D-galactopyranoside (ONPG) was added to a final concentration of 10 mg/mL and β-Galactosidase activity was measured by OD_420_ using a plate reader (BioTek Cytation 1 Cell imaging reader, Agilent) β-Galactosidase activity units are defined as (µmol of ONP formed min^-1^) × 10^6^/(OD_600_ × ml of cell suspension) and are reported as mean ± standard deviation.

### rGRIL-Seq

RNA enrichment and sequencing were performed as described previously (78). Briefly, *Salmonella* strains were inoculated in NSLB medium with 15 µg/ml Gentamicin, 100 µg/ml Ampicillin, 0.2% glucose and grown overnight at 37°C on a tissue culture rotator at 50 rpm. These cultures were then diluted to OD_600_ 0.01 into 50 mL of HSLB with the Gentamicin and Ampicillin and grown at 37°C water bath at 200 rpm until OD_600_ 0.5. Then, the *hilA* 5’UTR mRNA was induced by adding 1mM IPTG (final concentration) and incubating for 1 hour. Next, the T4 RNA ligase was induced by adding 0.2% L-Arabinose (final concentration) and incubating for an additional 20 min. After incubation, 2 OD of cells were centrifuged, and the cell pellet was fast frozen in liquid nitrogen and stored at −80°C. The next day, total RNA was isolated by hot-phenol extraction followed by DNase treatment. Starting with a total of 15 µg of isolated RNA, chimeric RNAs were enriched using eight capture oligo probes corresponding to regions of the *hilA* 5’ UTR and Oligo-dT magnetic Beads as described (78). Pull-down products were then sent to the Roy J. Carver Biotechnology Center at the University of Illinois for sRNA library preparation and sequencing using NovaSeq SP 2 x 150 nt.

### In vitro and in vivo competition assays

The University of Illinois Institutional Animal Care and Use Committee (IACUC) reviewed and approved all animal work. Procedures were performed in our AAALAC-accredited facility in accordance with university and PHS guidelines under protocol 21197. BALB/c mice (Envigo, 6 to 8 weeks old) were inoculated orally or intraperitoneally (i.p.) with bacterial suspension containing a 1:1 mixture of *ΔinvR::cm tetR* strain and WT *tetR* strain. Briefly, strains were grown separately for 16 h in LB at 37°C then mixed at a 1:1 ratio. Then, cell mixtures were diluted to the appropriate concentration in 0.1 M phosphate-buffered saline (pH 8) to a final concentration of ~5 ×10^8^ per 200 µL for oral infection. For intraperitoneal infections, cell mixtures were diluted in 1x PBS to obtain ~10^3^ CFU per 200 µL final concentration. Before oral infection, food and water were withheld for 4 h, and then mice were inoculated with 200 µL of inoculum by oral gavage. After oral infection, food and water were replaced immediately. For intraperitoneal infections, mice were inoculated with 200 µL of cell suspension by intraperitoneal (i.p.) injection. All inocula were diluted and plated on the LB tetracycline plates, and then replica plated on the chloramphenicol/tetracycline plates to determine the exact ratio of strains. After 3.5 days of infection, mice were sacrificed by CO_2_ asphyxiation followed by cervical dislocation and the spleens and distal small intestines were harvested from orally infected mice, while the spleens were collected from i.p. infected mice. Tissues were homogenized and serial dilutions were plated on LB containing tetracycline. After incubation, colonies were replica plated on chloramphenicol/tetracycline plates to determine the ratio of strains recovered. In vitro competition assays were conducted by subculturing 10^3^ CFU of the same inoculum used for the in vivo experiments into 5 mL of LB. The cultures were incubated for 16 h at 37°C with aeration. Then the overnight cultures were diluted and plated as above. The resulting colonies were replica plated onto chloramphenicol/tetracycline plates plates. The competitive index was calculated as (percentage of strain A recovered/percentage of strain B recovered)/(percentage of strain A inoculated/percentage of strain B inoculated). Student’s t-test was used for statistical analysis.

### Ethics statement

All animal procedures were reviewed and approved by the University of Illinois Institutional Animal Care and Use Committee. Experiments were conducted in an AAALAC-accredited facility following university and U.S. Public Health Service guidelines under protocol 21197. Every effort was made to minimize animal suffering.

## ACKNOWLEDGMENTS

This work was supported by the National Institutes of Health R35 GM139557 awarded to C.K.V., and R21 AI166495 awarded to J.M.S. We extend our gratitude to both current and former members of the Vanderpool and Slauch laboratories for supplying strains, plasmids, valuable advice, and insightful discussions.

---

**Figure S1A.**
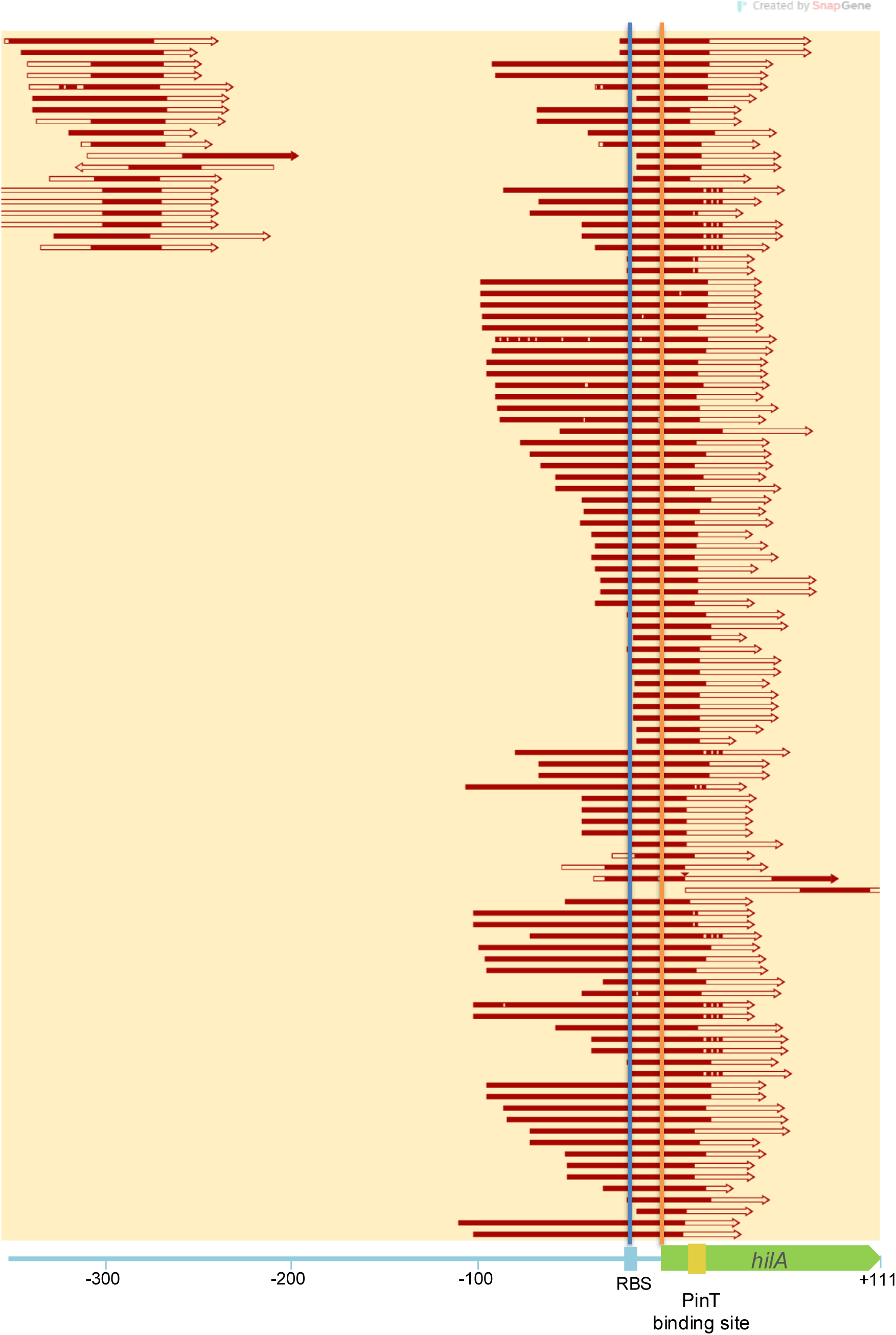
rGRIL-Seq captured the direct interaction between PinT and *hilA* mRNA 5’UTR. Schematic diagram of all PinT-*hilA* chimeric reads aligned to *hilA* mRNA. Blue vertical line denotes the ribosome binding site of *hilA* mRNA. Orange vertical line denotes the start codon of *hilA* mRNA. Red solid fragment indicates *hilA* fragment in chimeric reads, red outline arrow denotes PinT fragment in chimeric reads.

**Figure S1B.**
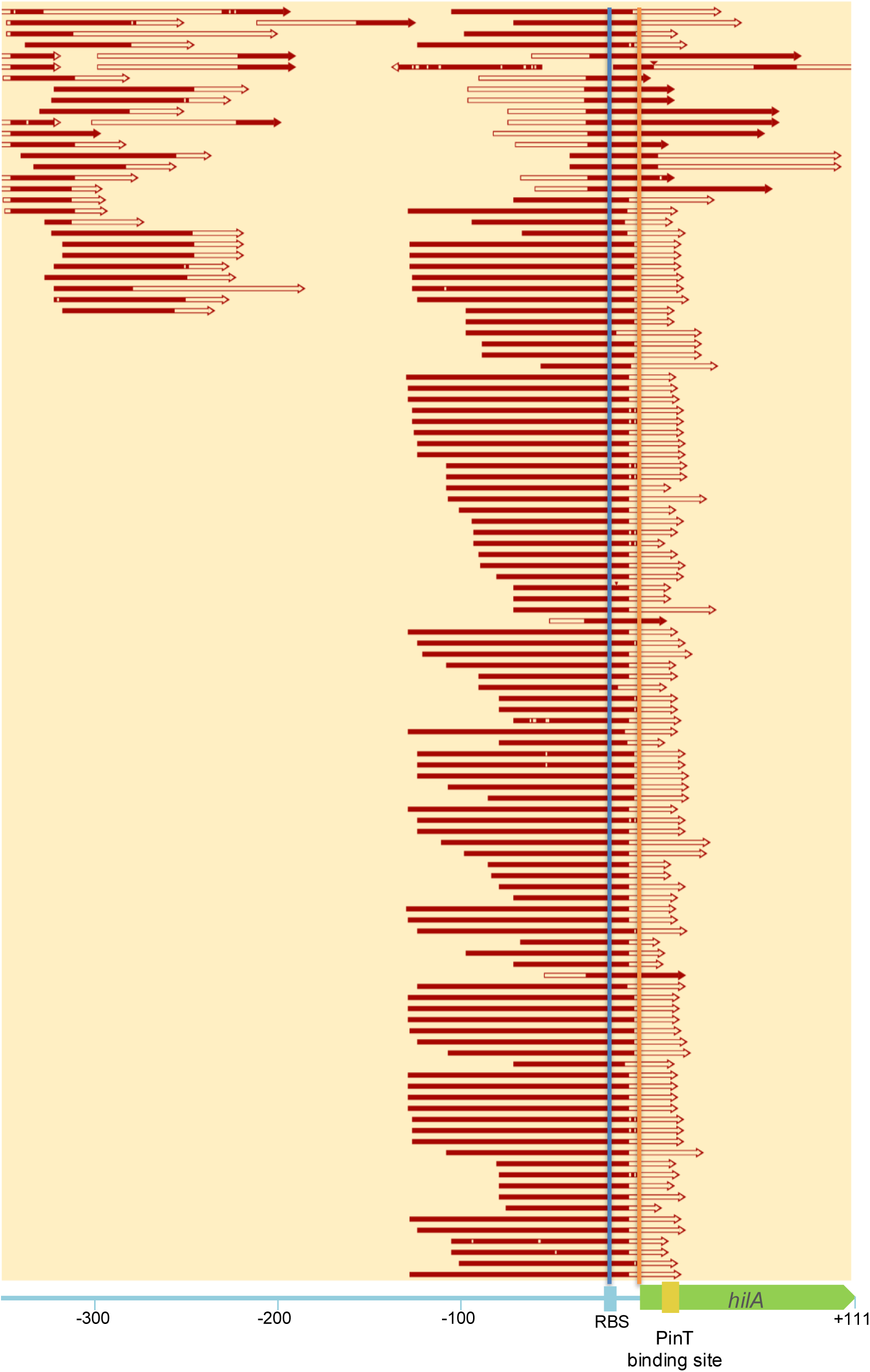
rGRIL-Seq captured the direct interaction between InvR and *hilA* mRNA 5’UTR. Schematic diagram of all InvR-*hilA* chimera reads aligned to *hilA* mRNA. Blue vertical line denotes the RBS of *hilA* mRNA. Orange vertical line denotes the start codon of *hilA* mRNA. Red solid fragment indicate *hilA* fragment in chimeric reads, red outline arrow denotes InvR fragment in chimeric reads.

